# EXPRESSED EMOTION AND SELECTED PATIENTS’ CLINICAL FACTORS AMONG CAREGIVERS OF SCHIZOPHRENIC PATIENTS VISITING JIMMA UNIVERSITY MEDICAL CENTER PSYCHIATRY OUT PATIENT UNIT, SOUTH WEST ETHIOPIA

**DOI:** 10.1101/2020.11.16.384396

**Authors:** Bethlehem Yimam, Matiwos Soboka, Yemiamrew Getachew, Bezaye Alemu, Gutema Ahmed, Elias Tesfaye

## Abstract

**Background:** Expressed emotion (EE) measures the emotion of the caregivers of persons with schizophrinia and is predictive of symptom levels in a range of medical and psychiatric conditions. It is worth to assess expressed emotion and associated factors among caregivers of patient with schizophrenia in Ethiopia since there is limited data on this issue in this part of the world.

**Objective:** To assess the status of expressed emotions and selecte patients’ clinical factors among care givers of patients with schizophrenia attending psychiatry oupatient unit of Jimma university medical center, South west, Ethiopia, 2019.

**Method:** A cross-sectional study design employed involving 422 caregivers of schizophrenic patients using consecutive sampling technique. Data was collected using structured interviewer administrated questionnaires (Family Questioners) which assess the level of expressed emotion, entered into Epidata 4.4 and analyzed by Statistical package for social science (SPSS) version 25. Descripitive statistics used to summerize data, bivariate logistic regression was done to identify candidate variables for multivariable logistic regressions and the association between expressed emotion and predictor variables was identified by using multiple logistic regression model.

**Results:** High expressed emotion was observed in 43.6% of respondents. Caring for schizophrenic patients for about 6-8 years, having 3-4 episodes of the illness was significantly associated with high expressed emotion.

**Conclusions:** This study revealed that there is high status of care givers expressed emotion compared to other studies. It also showed that number of episode of illnesses had significant association with high caregivers expressed emotion. Health care systems, which provide interventions for patients with schizophrenia, need to design proper strategy to address caregivers need as well.

## 1. INTRODUCTION

Expressed emotion measures the emotion of the care givers and is predictive of symptom levels in a range of medical and psychiatric conditions(1). Expressed emotion (EE) is an attitude, feeling, or behavior of the family caregiver in response to and reaction towards the person with schizophrenia(2).

Expressed emotion classification of caregivers is based mainly on the two variables ‘criticism’ (critical comments), and emotional over involvement, a third variable, ‘hostility’, is normally associated with high levels of critical comments. Those caregivers who showed high criticism or over involvement are rated as ‘high EE’ (9,12,13).

Schizophrenia is one of the most common serious mental disorders that result in changes in perception, emotion, cognition, thinking, and behavior. Both patients and their families often suffer from poor care and social isolation because of widespread ignorance about the disorder. In families with high levels of expressed emotion, the relapse rate for schizophrenia is high(6).

Approximately 50% of patients living with a spouse or their parents had at least one instance of readmission following discharge, compared with only 30% of those living alone(7).

Additional care giving role to already existing family roles becomes stressful psychologically as well as economically(7). Unemployment of both patients and families is a major indirect cost, resulting in more than half (61%) of the total economic burden of schizophrenia, these experiences lead family caregivers to have high expressed emotion (HEE), which in turn increases the risk of relapse of the person they are caring for(2).

A prospective study done in Brazil showed that 31% of patients presented relapses and, among the relatives, 68% presented elevated levels of expressed emotion. The proportions of family members with high levels of critical comments and emotional over involvement were 49% and 52%, respectively(3).

A study conducted in Nigeria showed the prevalence of high expressed emotion was 50.0% Relapse rates of people in differing living arrangements after an episode of mental disorder, and found that relapse rates were 17% for patients living alone or with siblings, 32% for those living with parents and 50%, for those living with spouse(8).

Patients with schizophrenia living with relatives who have a high expressed emotion (EE) level at admission to hospital are more likely to relapse within nine months after discharge than those patients whose relatives have a low EE level(9).

A cross sectional study conducted in Delhi, India schizophrenia patients having families with high level of critical comments had a three fold grater rate of relapse within 9 months after recovery and patients with high criticism have a larger chance of early relapse (10).

In psychiatric inpatient department of government medical College and hospital Nagpur the total mean score of the caregivers of patients with mental illness was 58.12 which shows high expressed emotions among caregivers, all the demographic factor results thecalculated value is less than tabulated value, so there is no association found between selected demographic variables and EE of the caregivers (11).

In the study conducted at outpatient clinics in Abbasia and Banha Hospitals for Mental Health it was found that statistically significant relationships existed between patients’ genders and parent EE; it was reported that parents of females made more critical comments than parents of males. Patients with adolescent onset more than half had parents rated high criticism(12).

The educational status of the demographic characteristic of patients and relatives was also significantly associated with high EE(13).

Hospital based cross sectional study conducted in India among 125 patients revealed that younger patient experienced more EE and Patients, who were single, experienced significantly more EE than married persons, which was similar with study done in Pakistan (14).

A study done in Nigeria showed Female care givers were associated with high expressed emotion. It has been found that younger age, female sex, higher educational level, and part-time occupation result into higher levels of psychological distressand distressed caregivers have high expressed emotion(18,26).

The British studies indicated that, among patients living in “high-EE” homes, the risk of relapse more than doubled for patients who were in face-to-face contact with relatives having high EE 35 hours per week or more (69% relapse rate) compared with those (28%) fewer than 35 weekly contact hours (16).

## 1. MATERIALS AND METHODS

### Study area and period

The study was conducted from April to June 2019 at Jimma University Medical Center (JUMC), which is found in Jimma town, south west Ethiopia

### Study Design

Institutional based cross sectional study design was employed.

### Population

All caregivers of patients with schizophrenia, visiting psychiatry out patient unit at Jimma university medical center were considered as source population. Caregivers who were ≥ 18 years of age and were taking care of patients with schizophrenia were included.

### Sample size determination and sampling technique

The Sample size was determined using single population proportion formula by taking the result done in Nigeria; the result of high expressed emotions of caregiver was 50.0%. To get the possible sample at 95% CI that is Z –value of 1.96 and marginal error of 5% is calculated as follow

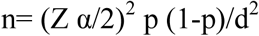

Where: n= number of sample size.

Z= desired 95% confidence, Z=1.96.

*p* = population proportion

q =1-p = 1-0.5=0.5

d *=* is the margin of sampling error tolerated (5%)

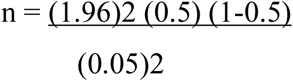

**n** initial =384

By considering 10% (10/100*384=38) non-response rate and final sample size was 422. Study participants wrre recruited by consecutive sampling techniques.

### Data Collection tool and Procedures

A structured questionnaire developed after reviewing related literatures was used to collect data about caregivers and patient socio-demographic variable and last psychiatric diagnosis was taken from medical records of patients.

The caregivers Expressed Emotion status was measured by Family Questioners (FQ) which is developed by Wiedemann, Rayki, Feinstein, and Hahlweg in 2002, with a 20 –items which include two domains– Critical Comments CC (10 items– 2, 4, 6, 8, 10, 12, 14, 16, 18, 20), with each maximum value 40 and the cut-off point for the FQ CC scale yielding maximum accuracy was a score of 23 (low<=23< high). Emotional Over Involvement EOI (10 items – 1, 3, 5, 7, 9, 11, 13, 15, 17, 19), with the maximum value 40 and the cut-off point yielding maximum accuracy was low <=27< high) of the relatives classified as high EOI. Possible responses are never or very rarely, rarely, frequently and very frequently, ranging from one to four, for each item. The FQ had better agreement with which is the gold standard questioners CFI on CC and EOI than did other short EE questionnaires, both have Sensitivity 80%, specificity 70%.Criticism (α=0.86, N=257) and emotional over involvement (α=0.80, N=256) subscales showed strong internal consistency(22,9, (18) 33,19,34).

### Study variable

#### Dependent variables

Expressed Emotion (EE) of caregivers

#### Independent variables

##### Care givers Socio-demographic factors

age, gender, ethnicity, occupation, marital Status, educational status, family size, relationship with patient, Distance from Hospital in km and average household monthly income.

##### Socio-demographic factors of patients

Age, gender, marital status,, educational status, Employment status, impact of illness on occupation

##### Clinical variables

duration of illness, duration of taking care of patient, time spent with the patient per day, number of episode, number of previous hospitalizations, number of family member with psychiatric illnesses, and co morbide disorder of the patients.

### Data processing, analysis and interpretation

The data was checked for consistency and completeness throughout the time of data collection. Data coded and entered twice in to double EPI-DATA version 4.41 and then exported to SPSS version 25 for analysis. Before performing binary regression the scores were checked for assumption and weather the model fit or not via Hosmerlemshow. Bivariate regression was computed for each independent variable separately with dependent variable. Finally those variables having p-value <0.25 taken to multiple logistic regressions model once and those variables with p-value of < 0.05on multiple logistic regression considered as having statistically significant association with the dependent variable.

### Ethical consideration

Ethical clearance was obtained from the Institutional Review Board of JU after approval of the proposal. Official permission was collected from Jimma university medical center psychiatry clinic. The purpose of the study was clearly communicated with study participants and data was collected after written consent is obtained. Caregivers with HEE were consulted to mental health professionals and psychologists working in the unit.

## 3. Results

### 3.1 Socio-demographic characteristics of study participants

A total of 422 caregivers of patients with schizophrenia were participated in this study. Among study participants, 281 (66.6%) were males, 263 (62.3%) were married, majority, 313 (74.2%) of respondents were Oromo by ethnicity and 313 (74.2%) were Muslims by religion. Mean age of participants was 40.24 years (SD ± 15.3) and 166 (39.3%) were parents. Nearly one-third (30.6%) of respondents attended primary education. Regarding occupation of the respondents, 146(34.6%) was farmer. More than half of the respondents, 228 (54%) live in urban areas, 110 (26.1%) live in distance of 9 - 23km from the Hospital and the median income was 1000 ETB (See table 1).

**Table 1:**
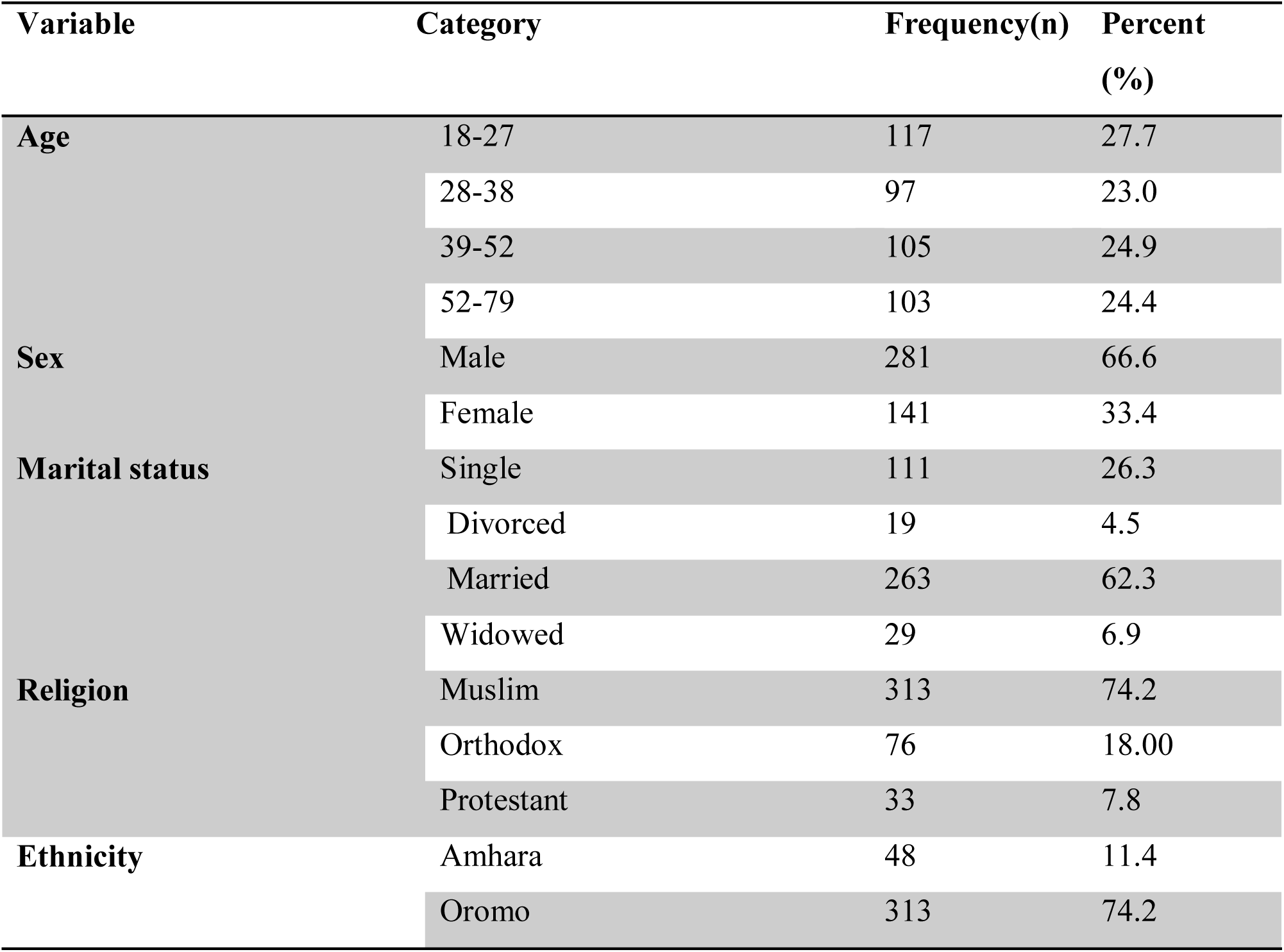
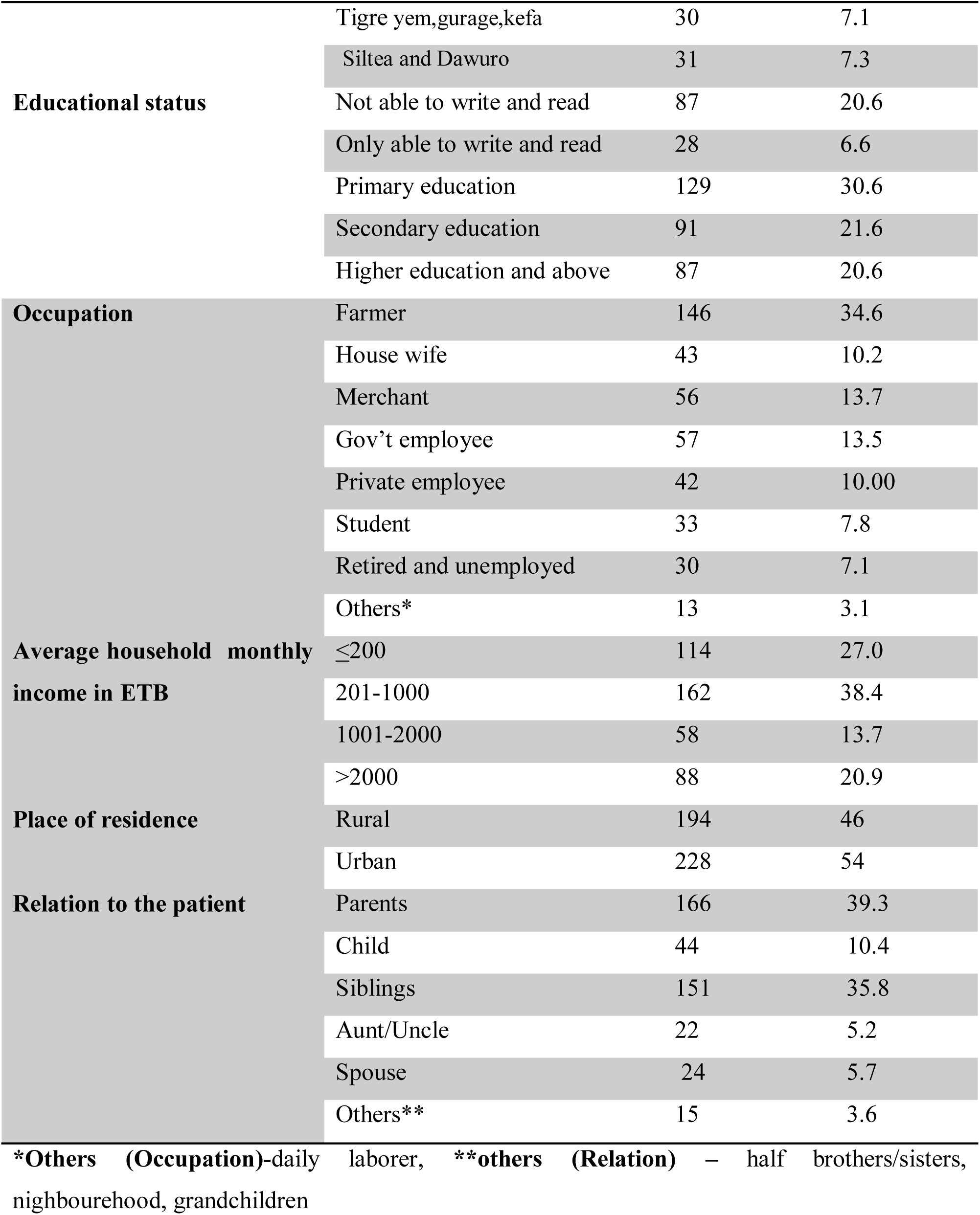
Socio-demographic characteristics of caregiver of patient with schizophrenia at JimmaUniversity medical center psychiatry clinic, South-west Ethiopia 2019 (n=422)

### 3.2 Caregivers perceived difficulties during care giving

From total respondants, 164(38.9%) had above seven family size. Three hundred seventy one (87.9%) had only one family member with mental illness. The mean duration of care giving was 5.7 (SD±4.18) years and the mean length of stay with the patient per twenty four hours was 7.49 (SD±6.24) hours. More than half of respondents, 260 (61.8%) had reported no objective burden and 173(41%) had reported sever subjective burden. Nearly all respondents had reported low perceived stigma 410(97.2%). Out of total respondents, 185(43.8%) had low social support. Nearly all of the participants 97.9% (n=413) reported as not having mental illness and 86.7% (n=366) not having chronic medical /physical illness which is reported by participants as diagnosed by health professional (See table2).

**Table 2.** Perceived difficulties during care giving and health status among caregiver of patient with schizophrenia at Jimma University medical center psychiatry clinic, South-west Ethiopia 2019 (n=422)

| Variable |  | Frequency(n) | Percent (%) |
| --- | --- | --- | --- |
| **Family size** | <=4family | 130 | 30.8 |
|  | 5-6 family | 128 | 30.3 |
|  | >7 family | 164 | 38.9 |
| **Family size with MI** | 1 family with MI | 371 | 87.9 |
|  | >2 family with MI | 51 | 12.1 |
| **Duration of take care of patient** | ≤2Years | 121 | 28.7 |
|  | 3-5 Years | 118 | 28.0 |
|  | 6-8 Years | 82 | 19.4 |
|  | >8 Years | 101 | 23.9 |
| **Relative's hours per day spent in contact with the patient** | ≤3hours | 129 | 30.6 |
|  | 4-6 hours | 126 | 29.9 |
|  | 7-12 hours | 117 | 27.7 |
|  | >12 hours | 50 | 11.8 |
| **Distance from Hospital in km** | ≤ 8 km | 106 | 25.1 |
|  | 9-23km | 110 | 26.1 |
|  | 24-50km | 107 | 25.4 |
|  | >50km | 99 | 23.5 |
| **Report of medical /physical illness** | Yes | 55 | 13.0 |
|  | No | 367 | 87.0 |
| **Report of mental disorder** | Yes | 8 | 1.9 |
|  | No | 414 | 98.1 |

### 3.3 Socio-demographic characteristics of the patients

The median age of the patient was 30 years and nearly one-third, 131 (31%) of the patients age was 25 and below. More than half, 310(73.5%) were males. Most of patients 271 (64.2%) were single and almost one fourth, 109 (25.8%) were married. One hundred eighty (42.7%) of patients attended primary education and 157 (37.2%) were unemployed. About 176(41.7%) patients had stopped their jobs due to the illness (See table3).

**Table 3:** Socio-demographic characteristics of patient with schizophrenia at Jimma University medical center psychiatry clinic South-west Ethiopia 2019 (n=422)

| **Variable** |  | **Frequency(n)** | **Percent (%)** |
| --- | --- | --- | --- |
| **Age** | 13-25years | 131 | 31.0 |
|  | 26-30years | 102 | 24.2 |
|  | 31-40years | 96 | 22.7 |
|  | 40-96years | 93 | 22.0 |
| **Sex** | Male | 310 | 73.5 |
|  | Female | 112 | 26.5 |
| **Marital status** | Single | 271 | 64.2 |
|  | Divorced | 30 | 7.1 |
|  | Married | 109 | 25.8 |
|  | Widowed | 12 | 2.8 |
| **Educational status** | Unable to read and Wright ,<br>only read and Wright | 89 | 21.1 |
|  | Primary education | 180 | 42.7 |
|  | Secondary education | 108 | 25.6 |
|  | Higher education and above | 45 | 10.7 |
| **Occupation** | Farmer | 101 | 23.9 |
|  | Housewife | 47 | 11.1 |
|  | Merchant | 18 | 4.3 |
|  | Gov't and Private Employee | 44 | 10.4 |
|  | Student | 38 | 9.0 |
|  | Unemployed | 157 | 37.2 |
|  | Others* | 17 | 4 |
| **Impact of the illness on occupational status** | Unemployed due to illness | 50 | 11.8 |
|  | Working full time | 77 | 18.2 |
|  | Working part time | 114 | 27.0 |
|  | Retired and Stop working | 181 | 42.9 |

\*Others, daily laborers

#### 3.3.1 Clinical characteristics of patient

Out of the total patients, 70 (16.6%) had co-morbid neuropsychiatric and medical disorder in summtion. Of this, 33 (7.8%) had substance use disorder as reviewed from their medical record. The mean duration of illness was 6.13 (SD±5.18) years and the mean age of first onset of illness was 26.28(SD±13.2) years. On the other hand, 295(69.9%) had 1-2 episodes and 259(61.4) of patients has no history of admission. (See table 4).

**Table 4:**
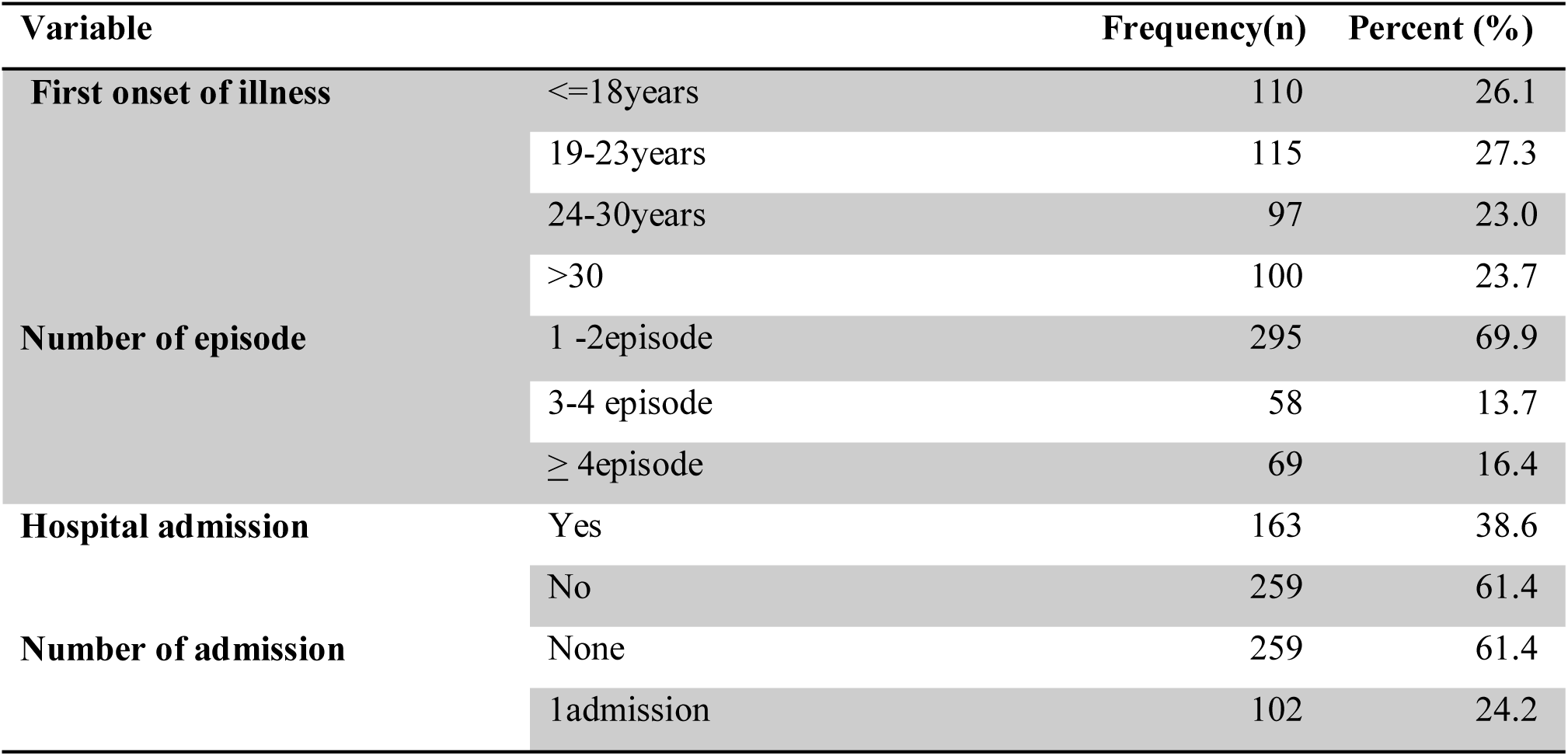
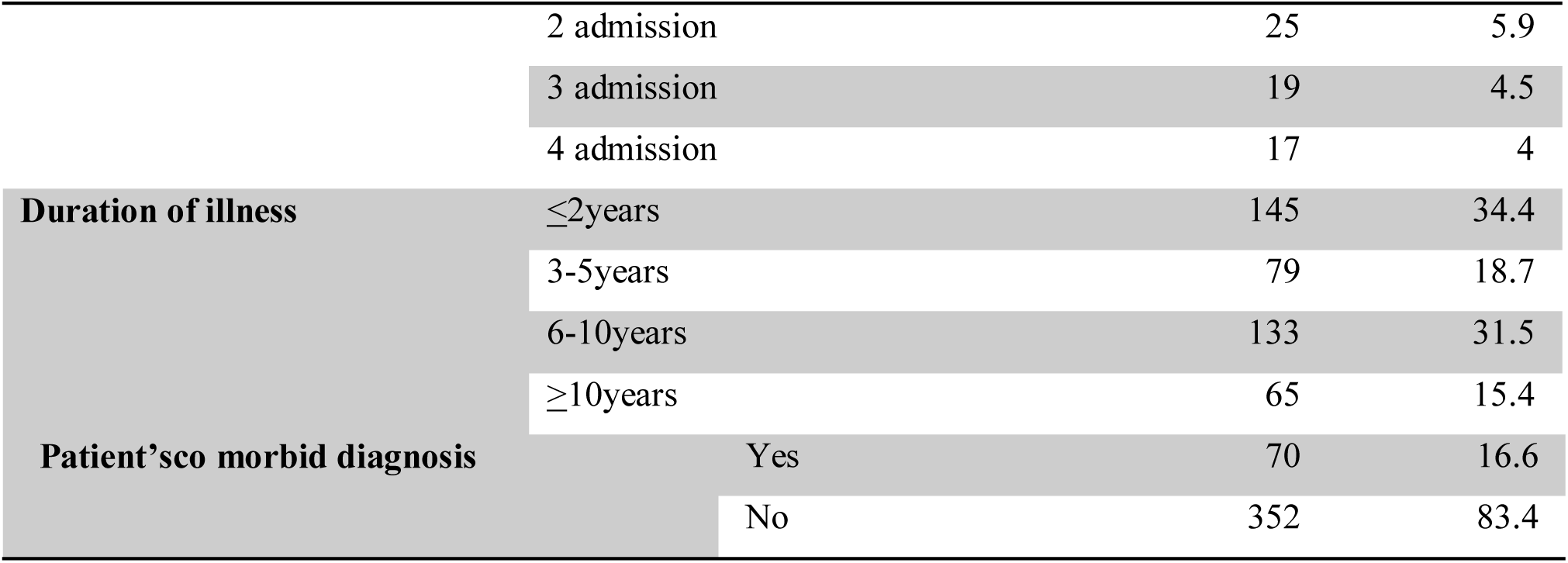
Clinical characteristics of patient with schizophrenia at Jimma University medical center psychiatry clinic, South-west Ethiopia 2019 (n=422)

### 3.4 Status of expressed emotions among caregivers of patient with schizophrenia

Of the total study participants, 101(23.9%) reported high critical comments (CC) and 148(35.1%) reported high emotional over involvement (EOI). Over all, the status of expressed emotion among caregivers as measured by considering either high CC or high EOI, 184[43.6% (38.5-48.6)] had higher expressed emotion (see Table 5).

**Table 5:** The family questioniers (FQ) sub scale among caregiver of patient with schizophrenia at Jimma University medical center psychiatry clinic, South-west Ethiopia 2019 (n=422)

| **Family Questioners(FQ) components for** |  | **Frequency(n)** | **Percent (%)** |
| --- | --- | --- | --- |
| **assessment of expressed emotion** |  |  |  |
| **Critical comments** | Low Critical Comment | 321 | 76.1 |
|  | High Critical Comment | 101 | 23.9 |
|  |  | 422 | 100.0 |
| **Emotional over involvement** | Low Emotional over involvement | 274 | 64.9 |
|  | High Emotional over involvement | 148 | 35.1 |
|  |  | 422 | 100.0 |
| **Expressed emotion status** | Low Expressed Emotion | 238 | 56.4 |
|  | High Expressed Emotion | 184 | 43.6 |
| **Total** |  | 422 | 100.0 |

### 3.5. Factors associated with expressed emotions among caregivers of patient with schizophrenia

#### 3.5.1. Bivariate analysis of factors associated with expressed emotion

Those who gavecare 6-8 years were found to be 2.4[COR=2.373, 95% CI (1.335, 4.218)], participants who were from a household with monthly income >2000 ETB were nearly 2 [COR=1.711, 95% CI((.976,2.999)], patients who had 3-4 episode were 2.3times [COR=2.382, 95% CI (1.339, 4.236)], (see in table 6).

**Table 6:**
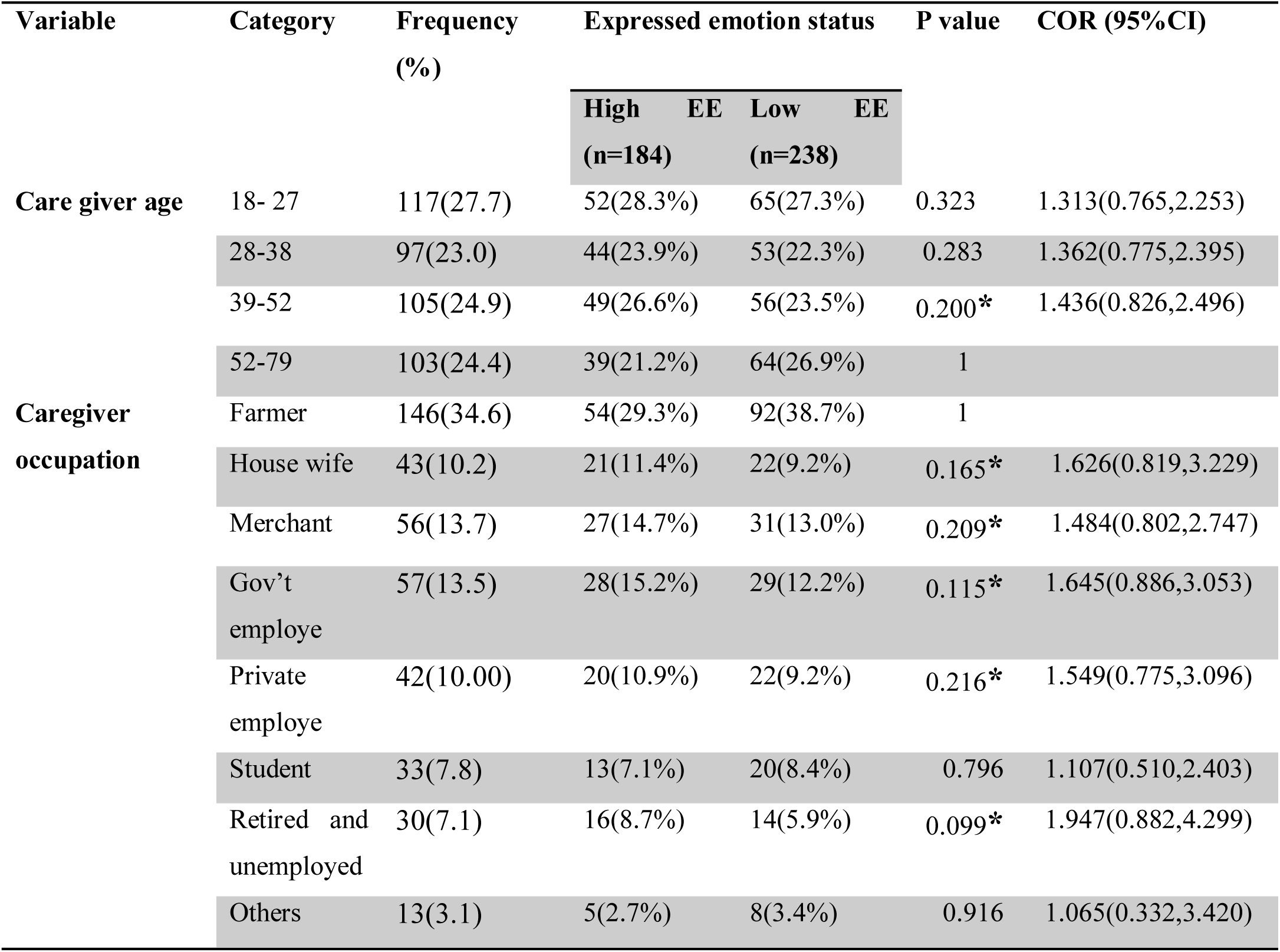
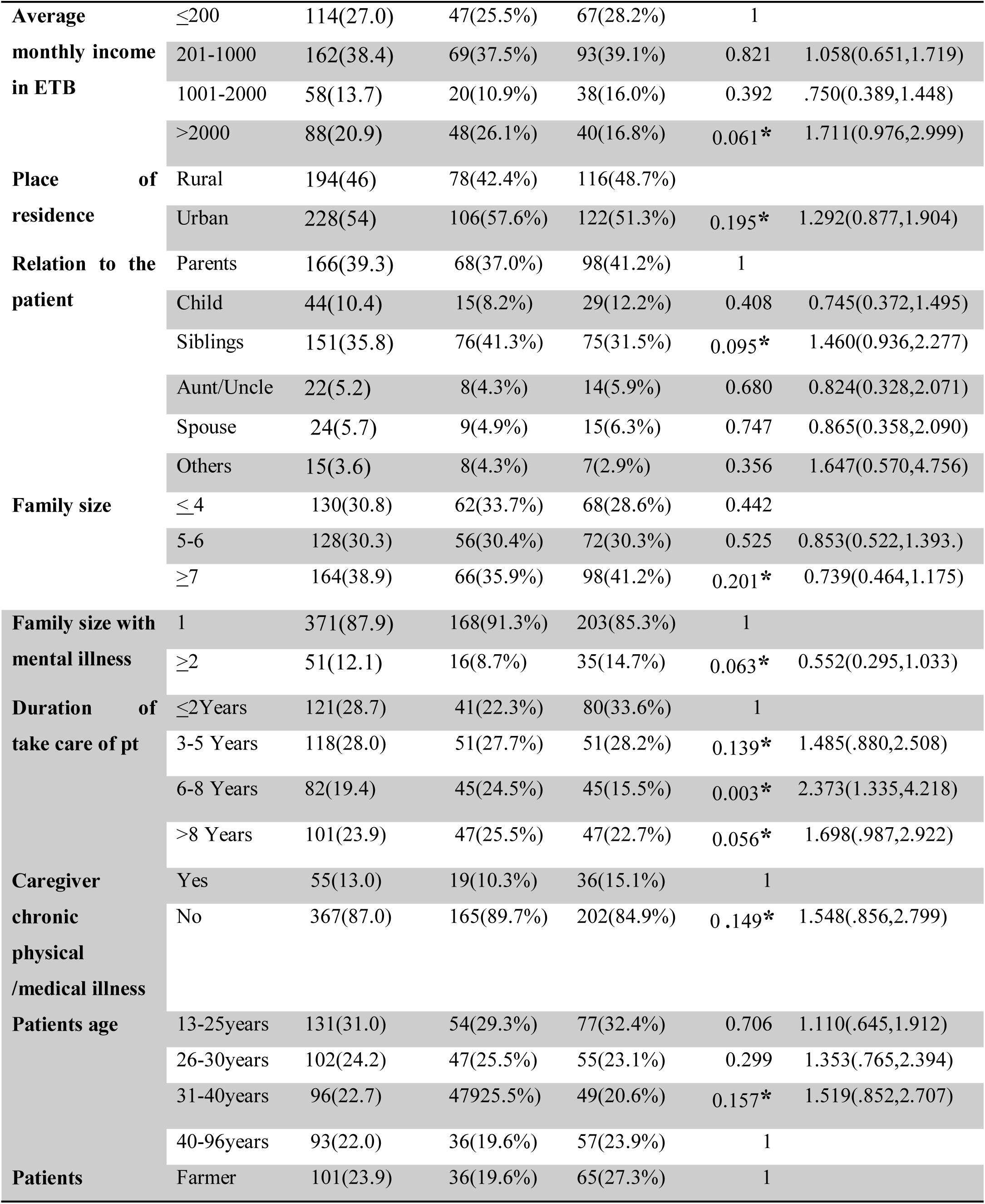
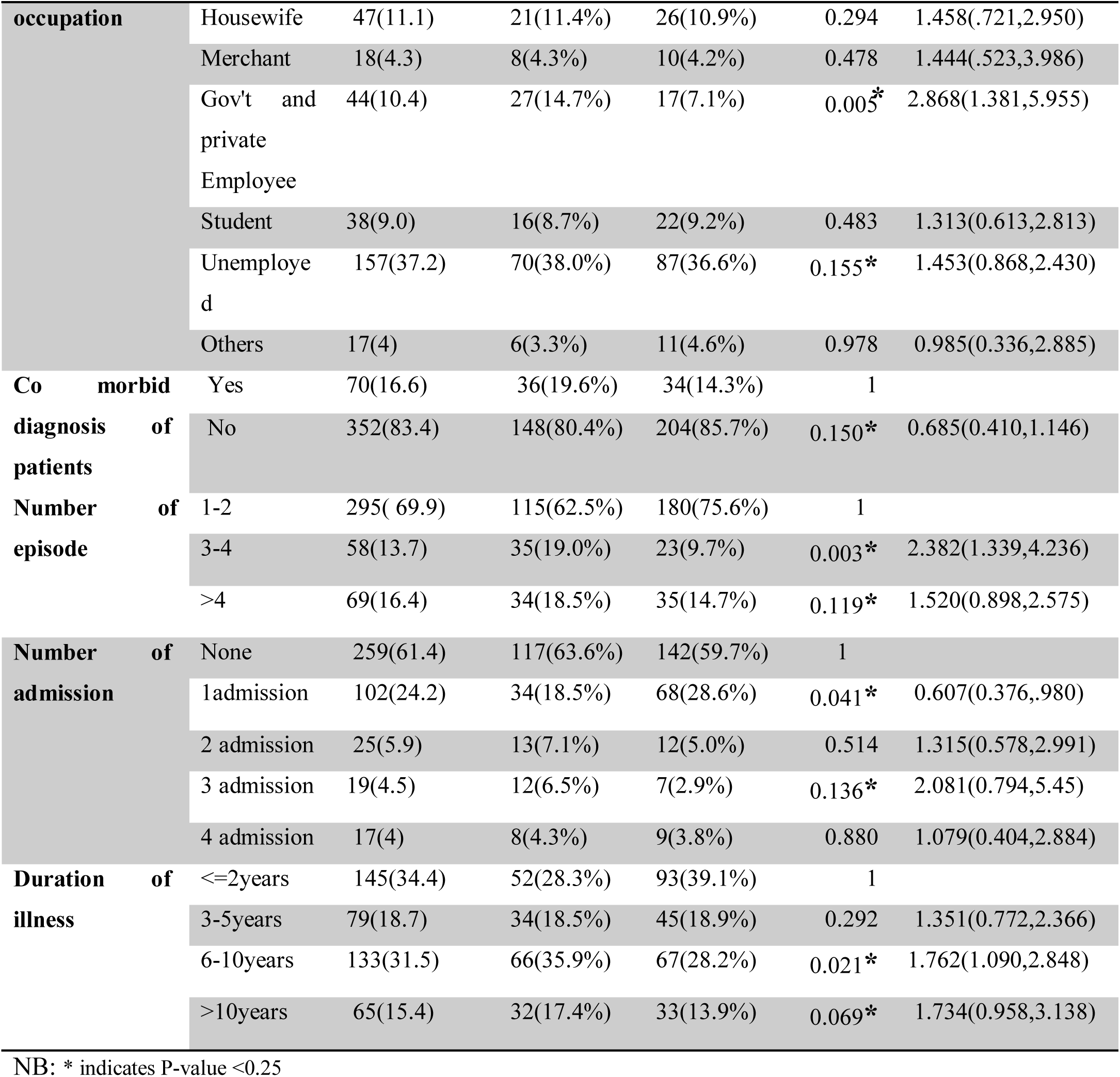
Bivariate analysis of factor associated and Status of expressed emotion among caregiver of patient with schizophrenia at JimmaUniversity medical center psychiatry clinic, South-west Ethiopia 2019 (n=422)

#### 3.5.2 Independent predictors of expressed emotions among caregivers of patient with schizophrenia at JUMC

Duration of giving care for 6-8 years were [AOR=2.439, 95% CI (1.308, 4.549)] caregivers report of no diagnosis of chronic medical/physical illness were [AOR=2.274, 95% CI (1.174, 4.406)] and patients who had 3-4 episode were 2.3times [AOR=2.281, 95% CI (1.253, 4.150)] were demonstrated to have statistically significant association with caregivers high expressed emotion.

The odds of having high expressed emotion among those who gave care for the patient for 6-8 years were 2.4 times higher than those who gave care < 2years.

The odds of having high expressed emotion 2.2 times higher than in those who had no chronic medical/physical illness than who had chronic medical/physical illness.

Finally, the odds of having high expressed emotion among those caregivers who had patients with 3-4 episode of illness were 2.3 times higher than in those who had 1-2episode of illness (See table 7).

**Table 7:**
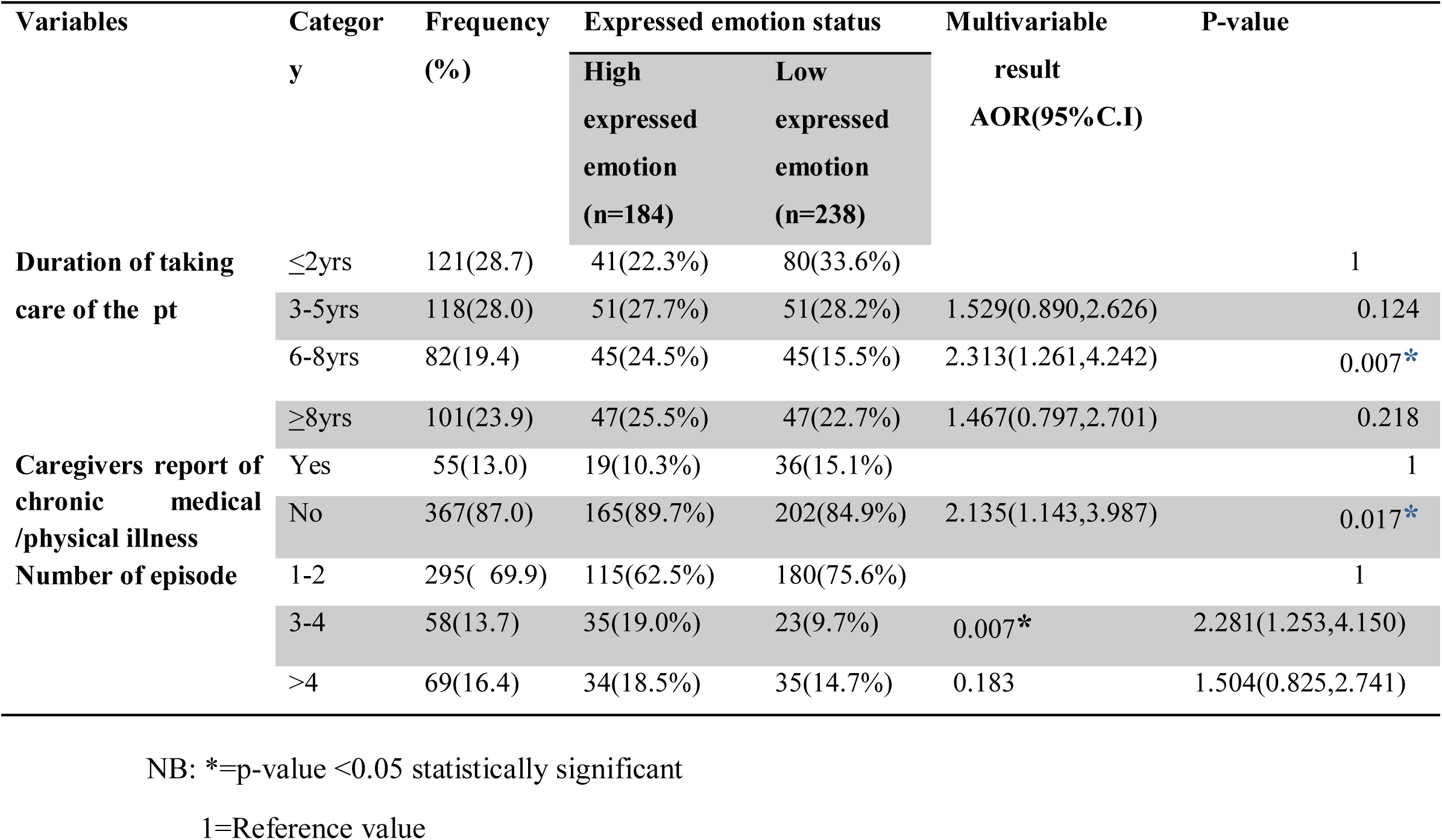
Multivariable logistic regression analysis of factors associated with high Expressed emotions among caregivers of patient with schizophrenia at Jimma University medical center psychiatry clinic, South-west Ethiopia 2019 (n=422)

| Variables | Category | Frequency (%) | Expressed emotion status |  | Multivariable result<br>AOR(95% C.I) | P-value |
| --- | --- | --- | --- | --- | --- | --- |
|  |  |  | High expressed emotion (n=184) | Low expressed emotion (n=238) |  |  |
| Duration of taking care of the pt | ≤2yrs | 121(28.7) | 41(22.3%) | 80(33.6%) |  | 1 |
|  | 3-5yrs | 118(28.0) | 51(27.7%) | 51(28.2%) | 1.529(0.890,2.626) | 0.124 |
|  | 6-8yrs | 82(19.4) | 45(24.5%) | 45(15.5%) | 2.313(1.261,4.242) | 0.007* |
|  | ≥8yrs | 101(23.9) | 47(25.5%) | 47(22.7%) | 1.467(0.797,2.701) | 0.218 |
| Caregivers report of chronic medical /physical illness | Yes | 55(13.0) | 19(10.3%) | 36(15.1%) |  | 1 |
|  | No | 367(87.0) | 165(89.7%) | 202(84.9%) | 2.135(1.143,3.987) | 0.017* |
| Number of episode | 1-2 | 295( 69.9) | 115(62.5%) | 180(75.6%) |  | 1 |
|  | 3-4 | 58(13.7) | 35(19.0%) | 23(9.7%) | 0.007* | 2.281(1.253,4.150) |
|  | >4 | 69(16.4) | 34(18.5%) | 35(14.7%) | 0.183 | 1.504(0.825,2.741) |

NB: \*=p-value <0.05 statistically significant
1=Reference value

## 4. Discussion

A total of 422 caregivers of patients with schizophrenia were included in this study. The proportion of high expressed emotion(EE) was 43.6 percent which is consistent with a similar studies conducted in Nigeria (41.4%) (21) and USA (43%). In our study high CC was 23.9% and 35.1% high EOI but the domains high CC and high EOI in USA were reported to be 19% and 50% respectively (3,5). The discrepancy may be due to used different assessment tool and sample size.

However, the prevalenc of EE found on our study was less than a study finding done Nigeria Lagos University (50%). The difference might be due to the size of sample involved in the study done in Nigeria was only 50 caregivers (8).

On the study done in Pakistan, 75% caregivers had high expressed emotion, which is almost two fold higher than the current study. The difference could be using different assessment tool, very much small sample size on the Pakistan study and the cultural difference between the two populations (22).

Study done in India reported that only 21% of caregivers of patients with schizophrenia had high expressed emotion (23) as compared to high (43.6%) expressed emotions in this study. The difference could be due to variation in sample size since only 100 caregivers were involved in India’s study but similarly gender, educational level, occupation, relationship with patient, contact per day with patient did not have significant association with level of EE as our study showed. Regarding the duration of taking care of the patient, those who give care for 6-8 years 82(19.4%) were 2.4 times more likely to have high expressed emotion than those who give care ≤ 2 years. Study done in Cairo showed caregivers do not practice any activities and hobbies; this may be because the caregivers spend a lot of time with the patient to provide care and the majority of caregivers were spending more than 12 care hours per day and this leads them to have HEE (12). The possible explanation for this might be because patients with schizophrenia may not be able to carry out daily activities by themselves and turn back to depend more on their caregivers. Consequently, family caregivers are likely to evaluate their life as being filled with interruptions. This belief of the caregivers about their own inability to manage severe symptoms might make them encounter repetitious long-term stress, causing them to have the reactions or behaviors found in the HEE. Similarly study showed in northern India the caregivers who showed sustained distress were likely to show high EE and have a longer caring history(9). In contrary, on one study done in India’s Assam hospital, the duration of care giving for patients didn’t have statistically significant association with high expressed emotion(23).

A participant who has no report of medical/physical illness diagnosis by health profesional was 2.2 times more likely to have high expressed emotion than who have no report of medical/physical illness of diagnosis by physician. This might be due to caregivers who has medical /physical illness diagnosed by health professional were responsible for having follow up program for the patients, helping through day to day activities since schizophrenic patient has difficulties in self helping behavior related to this caregivers might exhausted and have HEE than caregivers who have report of diagnosis of medical /physical illness.

Other might be because those caregivers who have medical/physical illness could not take the responsibility to taking care of the patient and spent more time with them since they have their own illness. This leads them to have short time contacting the patient, therfore they become less likely to have HEE.

Those caregivers of patients who had 3-4 episode of illness was 2.3 times more likely to have high expreesed emotion than those caregivers of patients who had 1-2episode of illness. Consistently with the current study a meta-analysis identified 27 articles reporting EE and psychiatric relapses in schizophrenia patients and confirmed that EE is a good predictor of schizophrenia relapses, especially in patients in the most chronic phase of the disease, current study result found no significant association between relapse and HEE (24). Inconterary with the current study, a prospective exploratory study done in Brazil showed relationship between psychiatric relapses and EEwas not demonstrated in a 24-month period. Expressed emotion was insufficient to predict relapses(3). However, most importantly, it needs standardized relapse instrument as well as further and deeper investigation to conclude about the link between expressed emotion and relapse rate

## ACKNOWLEDGEMENT

First, thank you God!

My sincere gratitude goes to Jimma University and Wollo University, my advisors Mr. Matiws Soboka, Dr. Bezaye Alemu, Mr. Gutema Ahmed and Dr. Elias Tesfaye. In addition, I value you as my teachers (Mr Yemiamrew) and mentors in the art of research (Mr Mamo). You all guided me through this study with patience and understanding. My gratitude also goes to my colleagues, (Workwa, Bennyam), the nurses and all other members of staff of the Department of Psychiatry

I am grateful to the all the caregivers of patients with schizophrenia who gave me the information which would help in this study and data collectors.

I love you mom and dad, you were my strength thank you.

